# Early blood transcriptomic markers of necrotizing enterocolitis in preterm pigs

**DOI:** 10.1101/2020.09.20.305631

**Authors:** Xiaoyu Pan, Tik Muk, Shuqiang Ren, Fei Gao, Per Sangild

**Affiliations:** Comparative Pediatrics and Nutrition, Department of Veterinary and Animal Sciences, Faculty of Health and Medical Sciences, University of Copenhagen, Copenhagen, Denmark; Genome Analysis Laboratory of the Ministry of Agriculture, Agricultural Genomics Institute at Shenzhen, Chinese Academy of Agricultural Sciences, Shenzhen, China

## Abstract

Preterm infants frequently develop necrotizing enterocolitis (NEC), a severe intestinal disorder associated with high mortality. Early detection of NEC is difficult due to poor specificity and sensitivity of clinical signs. We hypothesized that early development of NEC, before clear clinical symptoms appear, might affect expression of blood genes, potentially related to early systemic immune responses. Using preterm pigs as models for preterm infants, a retrospective analysis was performed on 129 infant formula fed pigs that had NEC diagnosis at necropsy on day 5. Clinical data including growth, activity, hematology, gastric residuals, incidence of diarrhea, bloody stool and abdominal distention were retrospectively reviewed. During this early postnatal period, except that bloody stool was observed in 19% of NEC pigs and absent in healthy pigs, no other clinical outcomes showed difference between NEC and healthy pigs. Whole blood transcriptome was compared between NEC pigs (NEC, n=20) and their litter-mate healthy controls (CON, n=19) on day 5, and revealed 344 differentially expressed genes (DEGs). PubMed literature search identified 123 genes that co-occurred with at least one of 9 NEC-related keywords (NEC, colitis, necrotic, hemorrhage, epithelial apoptosis, intestinal inflammation, inflammatory bowel disease, Crohn’s disease, ulcerative colitis). Co-expression network analysis suggested PAK2 as one of the hub genes. Using whole blood and dried blood spots (DBS) from another group of preterm pigs for validation, up-regulation of PAK2 and genes that co-occurred with NEC and other keywords in PubMed literatures (AOAH, ARG2, FKBP5 and STAT3) was confirmed in severe NEC cases. Specifically, *ex vivo* stimulation of cord blood with *S*.*epidermidis* increased ARG2. Our results show that whole blood gene expressions are affected in preterm pigs at an early stage when NEC is suspicious. Expression of target genes may be used to indicate NEC severity and associated bacterial infection. Routinely collected neonatal DBS may be used to develop early biomarkers for identifying infants with severe NEC lesions, thus providing better intervention strategy.

## Introduction

Every year, 15 million infants are born preterm (<37 weeks gestation). About 5-10% of preterm infants with very low birth weight (<1500g) are affected by necrotizing enterocolitis (NEC), a severe intestinal disease causing increasing deaths in preterm infants [1]. NEC develops rapidly and unpredictably during the first weeks after preterm birth and often coexists with sepsis. Currently, NEC diagnosis relies on a combination of clinical symptoms, including abdominal distention and bloody stools, and radiological features. Patients are initially treated with broad-spectrum antibiotics together with paused enteral feeding, and up to 50% of NEC cases require surgical intervention to remove necrotic intestine [2]. However, lack of sensitive and specific early diagnosis limits timely intervention against NEC, and 12-36% of patients die after NEC surgery [3, 4]. Therefore, biomarkers that can assist early diagnosis or predict the severity of NEC are needed and will help guide clinical decision-making.

Previous studies have identified some non-specific biomarkers, which are inflammation mediators, including acute phase proteins, cytokines, chemokines and cell surface antigens. The major drawback of these biomarkers would be their inability to differentiate NEC from sepsis or a proinflammatory state of the body [5]. Fecal inflammatory proteins (e.g. Calprotectin, S100A12) are more indicative of the site of injury, but representative fecal specimen is difficult to collect. Some proteins that are released by damaged intestinal tissue into bloodstream (e.g. L-FABP, I-FABP and TFF-3) have been found to increase in surgical NEC cases [6]. These may provide vital information for identifying infants who need close monitoring or early surgery, although they are unlikely to be useful for predicting early and mild NEC cases. A recent study showed that resistin-like molecule β (RELMβ), which is important for gut barrier function and regulates local immunity, was elevated in NEC cases as early as suspicious NEC occurred [7]. With recent advances in multi-omic technologies, investigators have focused on identifying novel and early biomarkers using these approaches. A study using microbiome and metabolome analysis has discovered high firmicutes dysbiosis and urinary alanine-to-histidine ratio as predictive biomarkers [8]. Another study profiling miRNA expression has identified plasma miR-1290 as specific biomarker which could differentiate NEC from neonatal sepsis or inflammatory conditions [9]. Adding novel information by using other omics techniques may help provide a more comprehensive picture of NEC biomarkers.

Numerous studies have been done to investigate the mechanism of NEC development. TLR4 activation has been suggested as a critical contributor to NEC [10]. It affects epithelial integrity by increasing enterocyte apoptosis and controlling enterocytes proliferation and migration [11], which may allow pathogen to enter circulation resulting in systemic inflammation. Moreover, TLR4 activation also mediates lymphocytes influx, leading to an induction of Th17 cells and reduction in Tregs. Release of inflammatory cytokine IL-17 by Th17 cells impairs enterocytes tight junction and causes mucosal injury during NEC development [12]. These results suggest a close relationship between NEC and immune response.

We hypothesized that early development of NEC might affect expression of blood genes, potentially related to early systemic immune responses. In this study, we aimed to characterize the whole blood transcriptome in preterm pig model of NEC and identify NEC-associated genes. Key genes were validated in both whole blood and dried blood spots to provide further clues for clinical use.

## Materials and Methods

### Animal experimental procedure

All animal procedures were approved by the Danish National Committee on Animal Experimentation. One hundred and seventy-three piglets from 8 sows (Danish Landrace x Large White x Duroc) were delivered by caesarean section at preterm (day 106, 90% of gestation). Immediately after caesarean section, all pigs were transferred to our piglet neonatal intensive care unit, fitted with orogastric and umbilical arterial catheters and reared in individual incubators. Preterm pigs received parenteral nutrition (2 to 4 mL/kg/h) plus a gradually increasing amount of enteral nutrition with infant formula (24 to 120 mL/kg/day) for 5 days to induce NEC. Pigs were weighed once daily and clinically assessed twice daily. Feces were scored according to following criteria: 1 = firm feces, 2 = pasty feces, 3 = droplets of watery feces/diarrhea, 4 = moderate amounts of diarrhea, 5 = large amounts of diarrhea. Presence of bloody diarrhea and/or abdominal distension was recorded. Physical activity of each pig was recorded by an infrared video surveillance camera and proportion of active time within each hour was recorded [13]. On day 5, intestinal permeability was measured by the lactulose-mannitol technique [14] shortly before euthanasia. Blood samples were collected by cardiac puncture at the time of euthanasia and immediately transferred into EDTA tubes for later hematology analysis. At necropsy, the entire small intestine was evenly divided into three regions (proximal, middle and distal intestine) and NEC was evaluated for both small intestine and colon according to a macroscopic NEC scoring system: 1 = absence of lesions, 2 = local hyperemia, 3 = hyperemia, extensive edema and local hemorrhage, 4 = extensive hemorrhage, 5 = local necrosis and/or pneumatosis intestinalis, 6 = extensive transmural necrosis and/or pneumatosis intestinalis. Pigs with a score ≥ 3 in any of the intestinal regions (including small intestine and colon) were diagnosed as “NEC”. During the study, 19 pigs died due to failure to mount independent respiration within the first 1-2 h after resuscitation attempts or later respiratory stress within 48 h. Twenty-five pigs were randomly selected to be euthanized on day 1 for other research propose. The remaining 129 pigs that survived until day 5 were used in this study for retrospective analysis of clinical data. Twenty NEC pigs (NEC) and 20 litter-mate healthy controls (CON) were randomly selected from 6 sows as discovery cohort and were used for transcriptome analysis. Forty-one pigs delivered from the other 2 sows were used as validation cohort to perform QPCR analysis.

### Transcriptome analysis

Blood samples from the discovery cohort were collected by cardiac puncture at euthanasia and 50 µl of whole blood was stored in Lysis/Binding solution (MagMax 96 blood RNA isolation kit, Thermo Fisher, USA) at -20 °C. Total blood RNA was extracted using MagMax 96 blood RNA isolation kit according to the manufacturer’s instructions, no more than 6 months after storage. RNA-seq libraries were constructed using 500 ng RNA and VAHTS mRNA-seq V3 Library Prep Kit for Illumina (Vazyme, China). The libraries were sequenced on the Illumina Hiseq X Ten platform (Illumina, USA) to generate 150 bp paired-end reads. Quality and adapter trimming of raw reads were performed using TrimGalore (Babraham Binoinformatics, UK). The remaining clean reads (∼ 26 M per sample) were aligned to the porcine genome (Sscrofa11.1) using Tophat2 [15]. The annotated gene information of porcine genome was obtained from Ensembl (release 91). The script htseq-count [16] was used to generate gene count matrix, followed by gene-level expression analyses using DESeq2 [17].

### QPCR analysis

Blood samples from the validation cohort were collected by cardiac puncture at euthanasia and stored by two methods for up to one year before analysis: 1) stored as whole blood (WB) in the Lysis/Binding solution as described above and 2) stored as dried blood spots (DBS). For both storage methods, 50 µl of whole blood was used. For the DBS sample, blood was spotted onto a filter paper (gift from Statens Serum Institut, Denmark) using a pipette, and was air-dried at room temperature for about 3 hours before storage at -20 °C. Total blood RNA from the 1^st^ storage method was extracted using MagMax 96 blood RNA isolation kit according to the manufacturer’s instructions. Total blood RNA from DBS was extracted using Monarch Total RNA Miniprep Kit (New England Biolabs, USA) with a modified protocol. Briefly, one full DBS was cut into small pieces and incubated in 600 µl 1X DNA/RNA Protection Reagent at room temperature. During incubation, the DBS sample was homogenized by continuous vortexing with a stainless steel bead (Sigma-Aldrich, # Z763829, USA) for 15 minutes, followed by Prot K reaction at 55 °C for 30 minutes. Supernatant was collected using Qiashredder (Qiagen, Germany) by centrifugation at 10000 g for 2 minutes and RNA was extracted following the rest of protocol.

Blood RNA after extraction was quantified using NanoDrop 1000 spectrophotometer (Thermo Fisher, USA). For WB and DBS samples, 500 and 50 ng RNA were converted into cDNA respectively using High-Capacity cDNA Reverse Transcription Kit (Thermo Fisher, USA). QPCR was performed using LightCycler 480 SYBR Green I Master kit (Roche, Switzerland) on LightCycler 480 (Roche, Switzerland) and results were analyzed according to double delta Ct method. Relative expression of target genes was normalized to the housekeeping gene *HPRT1*. All primers were designed using Primer-Blast [18] and the sequences of primers are listed in Supplementary Table S1.

### *Ex vivo* whole blood stimulation

Cord blood from 12 newborn preterm pigs was collected for *ex vivo* stimulation with live bacteria *S*.*epidermidis* (10^7^ cells/mL, WT-1457, received from Havard Medical school) and lipopolysaccharides (LPS, 1 µg/mL, E. coli 0111:B4, InvivoGen), respectively. Stimulation with *S*.*epidermidis* and LPS was performed at 37°C and 5% CO2 for 2 and 5 h, respectively. After stimulation, cord blood RNA was extracted using MagMax 96 blood RNA isolation kit and was used to perform QPCR as described above.

### Statistical analysis

For clinical outcomes, comparison between two groups (NEC vs. CON) was performed using t-test for continuous variables (birth weight, daily weight gain, physical activity, gut permeability, gastric residuals and hematology data), or Fisher’s exact test for independence for categorical variables (sex, presence of diarrhea, bloody stool and abdominal distension). A p-value <0.05 was considered statistically significant, and a p-value <0.10 was considered a tendency to an effect. For transcriptome analysis, significant differentially expressed genes (DEGs) between NEC and CON groups were identified by DESeq2 using Benjamini-Hochberg (BH)-adjusted p-value <0.1 as cut-off. Correlation between gene expression was performed using Spearman’s rank correlation based on normalized counts produced from DESeq2, and the correlation with absolute spearman rho >0.6 as well as BH-adjusted p-value <0.05 was considered statistically significant. Gene ontology and KEGG pathway enrichment analysis were performed using DAVID [19] and a BH-adjusted p-value <0.05 was considered statistically significant. For QPCR analysis, comparison between two groups was made using t-test and a p-value <0.05 was considered statistically significant.

## Results

### Clinical outcomes at early NEC stage

A total of 129 preterm pigs from 8 sows were euthanized on postnatal day 5 for NEC diagnosis as scheduled. Necropsy found 74 NEC cases and 55 healthy controls (CON). According to NEC severity (defined as the maximum NEC score among all intestinal regions), the 74 NEC pigs included 49 mild (score 3-4) and 25 severe cases (score 5-6). Colon NEC lesions were observed in 53 cases, while the other 21 NEC cases had small intestinal lesions with or without colon lesions. The location of NEC lesions was independent of NEC severity (P >0.05). Clinical outcomes of all 129 preterm pigs are summarized in Table 1.

**Table 1.** Clinical outcomes of preterm pigs.

|  | CON (n=55) | NEC (n=74) | Significance |
| --- | --- | --- | --- |
| Birth weight, g | 1003±32 | 1027±33 | ns |
| Daily weight gain, g/d | 13±2 | 14±2 | ns |
| Female, n (%) | 25 (45%) | 36 (51%) | ns |
| Diarrhea, n (%) | 41 (74.5%) | 59 (79.7%) | ns |
| Bloody stool, n (%) | 0 (0%) | 14 (18.9%) | *** |
| Abdominal distention, n (%) | 4 (7.3%) | 3 (4.1%) | ns |
| Active time per hour (%) | 16.60±0.86 | 15.48±0.61 | ns |
| Gut permeability, L/M ratio | 0.14±0.03 | 0.14±0.02 | ns |
| Gastric residuals, g/kg | 19.75±1.27 | 22.49±1.24 | ns |
| Hematology |  |  |  |
| Total leukocytes, 10<sup>9</sup>/L | 2.63±0.21 | 2.45±0.19 | ns |
| Total erythrocytes, 10<sup>12</sup>/L | 3.84±0.10 | 3.76±0.09 | ns |
| Hemoglobin, mmol/L | 5.18±0.14 | 5.05±0.12 | ns |
| Hematocrit, L/L | 0.28±0.01 | 0.28±0.01 | ns |
| MCV, fL | 72.98±0.63 | 71.64±0.49 | # |
| MCHC, mmol/L | 18.51±0.17 | 18.82±0.14 | ns |
| Platelets, 10<sup>9</sup>/L | 323.78±23.91 | 266.24±19.95 | # |
| MPV, fL | 13.37±0.41 | 12.92±0.31 | ns |
| MPC, g/L | 223.58±1.71 | 224.56±1.15 | ns |
| Neutrophil, % | 44.41±1.84 | 46.62±1.56 | ns |
| Lymphocyte, % | 50.71±1.86 | 48.74±1.54 | ns |
| Monocyte, % | 2.73±0.21 | 2.51±0.20 | ns |
| Eosinophil, % | 0.85±0.16 | 1.20±0.15 | ns |
| Basophil, % | 0.22±0.03 | 0.21±0.04 | ns |
| LUC, % | 1.07±0.15 | 0.80±0.08 | # |

Data are presented as means ± SEMs or numbers (\*P<0.05, \*\*P<0.01, \*\*\*P<0.001, #P<0.1). LUC, large unstained cells; MCHC, mean corpuscular hemoglobin concentration; MCV, mean corpuscular volume; MPC, mean platelet component; MPV, mean platelet volume.

The average birth weight of preterm pigs was 1017 g and was similar between NEC and CON groups (1027±33 vs. 1003±32 g, P >0.05). During the first 3 days, diarrhea (feces score > 2) was observed only in a few pigs (up to 6 per group). From day 4, 67 out of 129 pigs had diarrhea but the incidence was similar between the two groups (79.7 vs. 74.5% for NEC vs. CON, P >0.05). Bloody stool was observed in 14 NEC pigs but was absent in CON pigs (P <0.05). Of note, almost all of the pigs (12/14) that had bloody stool were later diagnosed as severe NEC and only 2 mild NEC cases had bloody stool. Moreover, only 3 NEC pigs and 4 CON pigs showed abdominal distention (P >0.05). By day 5, NEC and CON pigs showed similar growth rate (14±2 vs. 13±2 g/d, P >0.05) and physical activity as detected by surveillance camera (15.5 vs. 16.6% active time per hour, P >0.05). At the time of euthanasia, two groups of pigs had similar gut permeability, gastric residual volume and hematology results, except for a tendency of decreased mean corpuscular volume (MCV), decreased platelets and decreased large unstained cells (LUC) in the NEC group (all P <0.1). In summary, during the early postnatal period, except that 19% of NEC pigs (especially severe NEC cases) showed bloody stool, no other clear clinical symptoms occurred before NEC diagnosis at necropsy.

### Blood transcriptome reveals NEC-associated differentially expressed genes (DEGs)

To examine whether whole blood gene expression changes at early NEC stage, blood samples were collected at euthanasia and 40 pigs from 6 sows were randomly selected for RNA-seq as discovery cohort. One sample failed in RNA-seq, resulting in 39 transcriptome data (20 for NEC and 19 for CON). Of 25,880 genes annotated in the Ensembl database (Sscrofa11.1, release 91), 18,894 genes were detected in at least one pig. Based on the individual expression of these genes, principal component analysis (PCA) showed no clear separation between NEC and CON groups (Figure 1A). To identify DEGs between the two groups, a threshold of adjusted *P* < 0.1 was used. This resulted in 344 DEGs with ascertainable gene symbols, including 158 and 186 DEGs that were up- and down-regulated in NEC, and equivalent to 1.8% of analyzed genes (Figure 1B). These results suggested that whole blood gene expression was not massively affected at early NEC stage. We further took a detailed look at the relationship between DEGs and NEC severity. The 20 NEC cases in our discovery cohort included 11 mild and 9 severe NEC cases. By comparing these two subgroups to CON pigs, we found that only 8 out of the 344 DEGs showed significant difference between mild NEC and CON groups, while majority of the DEGs (70%) were significantly altered by severe NEC. Therefore, at early NEC stage, similar to the clinical outcomes whereby mild NEC cases were hard to be distinguished from the healthy ones, change of whole blood gene expression was mainly driven by severe NEC.

**Figure 1.**
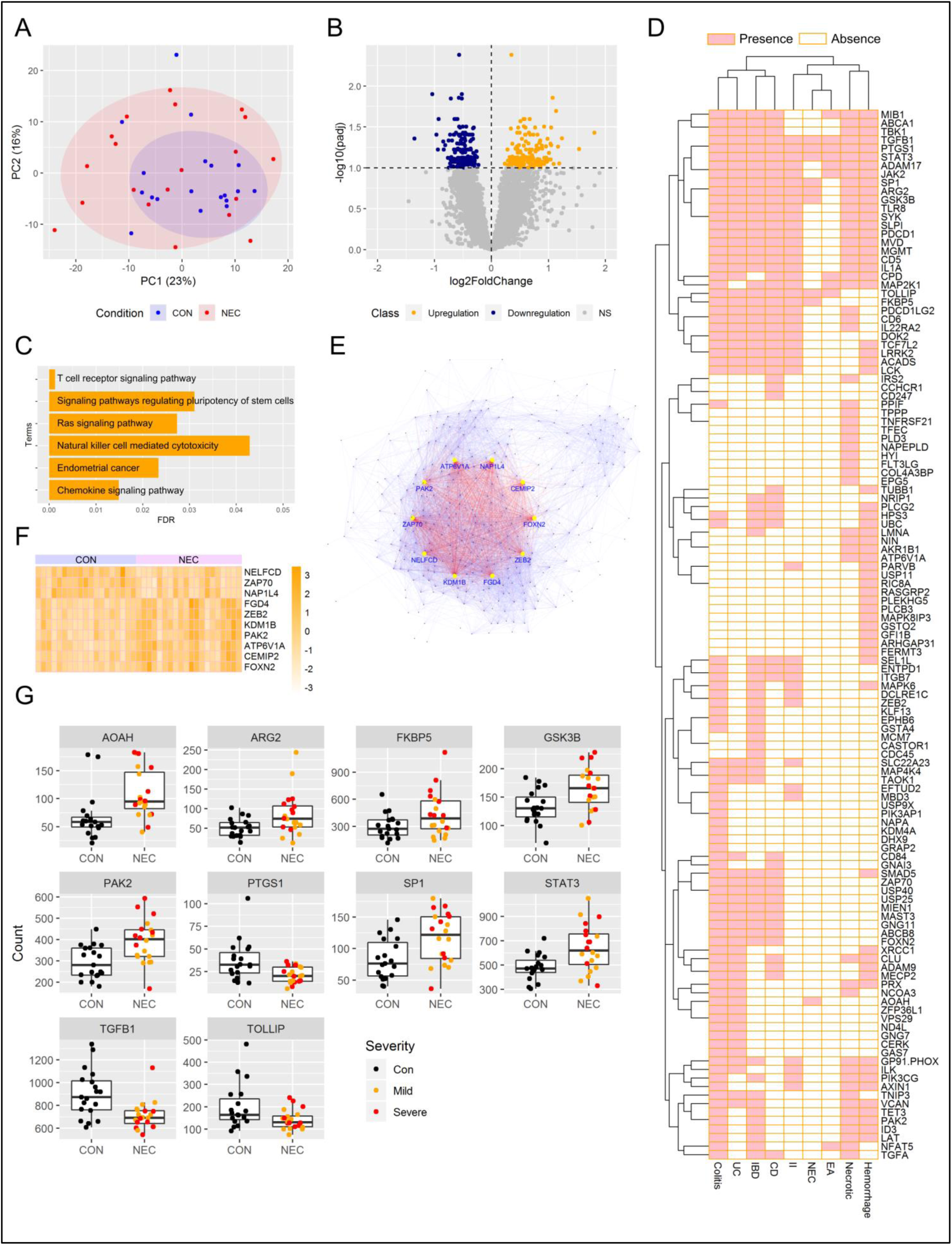
Whole blood transcriptome in response to NEC. (A) Principal component analysis (PCA) was performed based on the individual expression level of all analyzed genes. Scores of first two principle components from PCA are plotted. (B) Volcano plot illustrating the DEGs between NEC and CON group. (C) Barplot showing significantly enriched KEGG pathways. (D) Heatmap illustrating co-occurrence of DEGs and NEC-related keywords in PubMed literatures. Pink and white colors indicate for presence and absence of co-occurrence, respectively. (E) Putative co-expression network, where each node represents a gene and each edge represents a significant correlation between gene pairs. Top 10 hub genes are labeled in blue, and their edges are labeled in red. (F) Heatmap showing relative expression level of the top 10 hub genes between NEC and CON groups. (G) Boxplot of targeted DEGs that were selected for validation.

To understand the functional significance of these NEC-associated DEGs, KEGG pathway enrichment analysis was performed according to DAVID database. This resulted in 6 significant pathways (Figure 1C, FDR <0.05), including “T cell receptor signaling pathway”, “Chemokine signaling pathway”, “Endometrial cancer”, “Ras signaling pathway”, “Signaling pathways regulating pluripotency of stem cells” and “Natural killer cell mediated cytotoxicity”. Next, to explore whether any of the DEGs might play a role in NEC development, a subsequent PubMed literature search was performed. We defined 10 NEC-related keywords, including NEC, colitis, necrotic, hemorrhage, pneumatosis, epithelial apoptosis, intestinal inflammation, inflammatory bowel disease (IBD), Crohn’s disease and ulcerative colitis. Then for each DEG, we searched PubMed record using the “Text Word” field tag to identify co-occurrence of keywords and genes. Result showed no DEGs co-occurred with pneumatosis but revealed 123 DEGs that co-occurred with at least one of the other 9 keywords (Figure 1D). Moreover, we sought to characterize the relationships among all DEGs. To do this, pairwise correlation was performed to determine possible co-expressed gene pairs. A total of 6,830 out of 58,996 gene pairs having absolute Spearman rho >0.6 and FDR <0.05 were selected to conduct a putative co-expression gene network (Figure 1E). Among the top 10 hub genes (Figure 1F) that had the highest number of connection to other genes, *ATP6V1A, FOXN2, PAK2, ZAP70* and *ZEB2* were found in PubMed record and related to hemorrhage, IBD or colitis [20-24]. Finally, 10 DEGs including *PAK2* and 9 DEGs (*AOAH, ARG2, FKBP5, GSK3B, PTGS1, SP1, STAT3, TGFB1* and *TOLLIP*) that co-occurred with NEC together with other keywords in PubMed literatures were selected for validation (Figure 1G).

### Validation of NEC-associated DEGs in whole blood and dried blood spots

To validate the selected NEC-associated DEGs, 41 pigs delivered from 2 separate sows were used as validation cohort. This included 13 CON and 28 NEC pigs (incl. 22 mild and 6 severe NEC cases). We first examined the expression of 10 selected DEGs in whole blood by using QPCR. As a result, 5 genes (*AOAH, ARG2, FKBP5, PAK2* and *STAT3*) were increased in severe NEC cases, as compared to mild NEC cases or CON group (Figure 2A). However, when combining both mild and severe NEC cases as one NEC group, none of the selected DEGs showed significant differential expression between NEC and CON groups.

**Figure 2.**
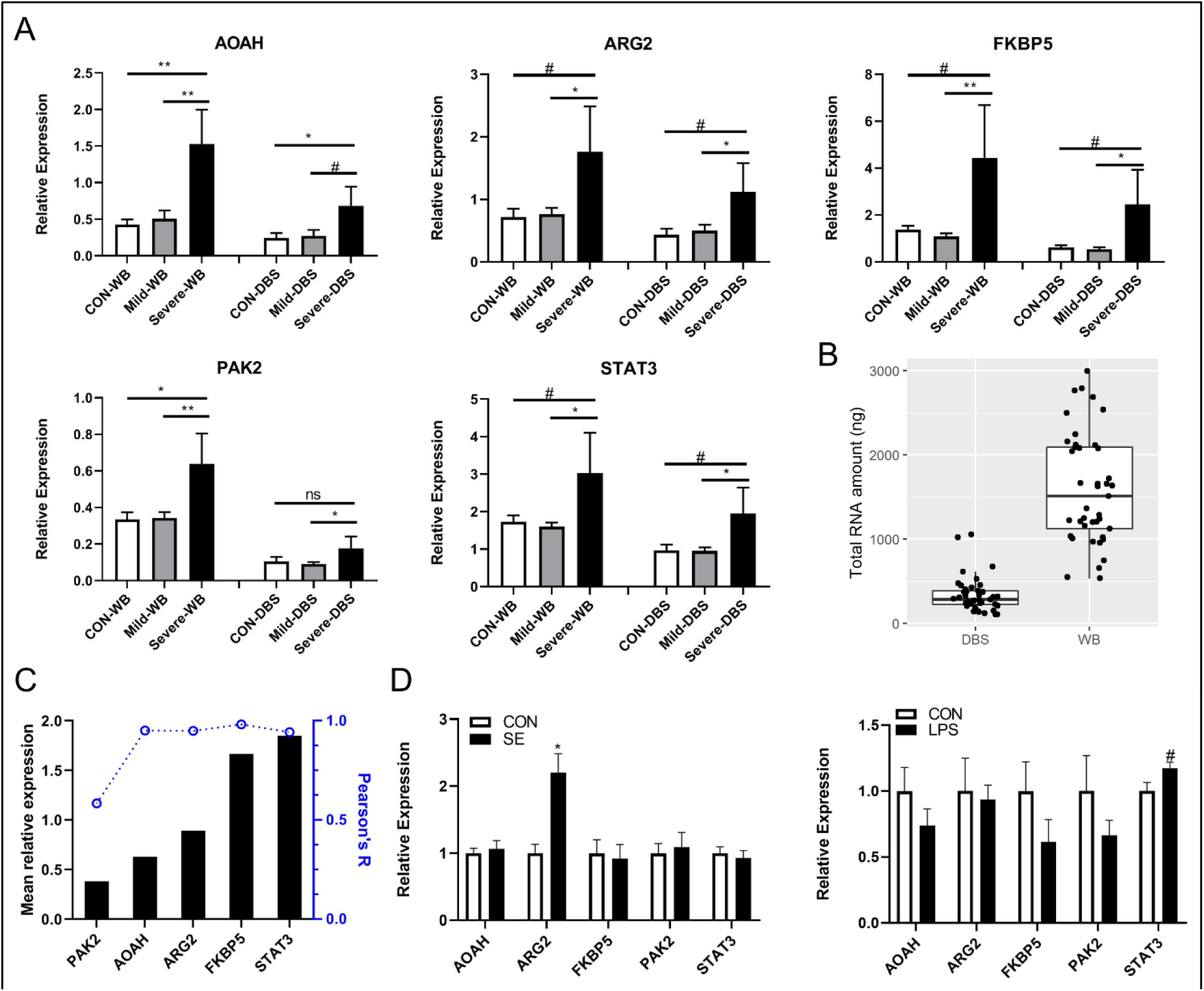
Expression of target DEGs in validation cohort and after *ex vivo* stimulation. (A) Expression of target DEGs in validation cohort. Animals were classified into 3 groups according to NEC score: CON (score 1-2, n=13), Mild (score 3-4, n=22) and Severe (score 5-6, n=6). Data are presented as means ± SEMs. Comparison between Mild vs. CON group or Severe vs. CON group was performed using t-test (*P<0.05, **P<0.01, ***P<0.001, #P<0.1). WB, whole blood; DBS, dried blood spots. (B) Boxplot showing total RNA amount extracted from 50 µl of whole blood (WB) or dried blood spots (DBS). (C) Correlation of gene expression between whole blood and dried blood spots. (D) Expression of target genes in cord blood after *ex vivo* stimulation with *S*.*epidermidis* (SE) and LPS. Data are presented as means ± SEMs. Comparison between treatment vs. control group was performed using paired t-test (*P<0.05, **P<0.01, ***P<0.001, #P<0.1).

As dried blood spot (DBS) samples are routinely collected as a part of the neonatal screening program worldwide, we next evaluated whether the expression pattern of these NEC-associated DEGs remained detectable in DBS, thus providing evidence for developing NEC biomarker using archived infant DBS. We found that an average of 1.6 µg total RNA could be extracted using 50 µl of whole blood, while DBS containing the same amount of blood revealed an average of 336 ng total RNA, equivalent to 21% RNA amount in the whole blood (Figure 2B). Nevertheless, QPCR quantification showed that the 5 target genes (*AOAH, ARG2, FKBP5, PAK2* and *STAT3*) in DBS were consistently increased in severe NEC cases (Figure 2A) and the expression levels of these genes in DBS significantly correlated with that in whole blood (Figure 2C). According to Pearson’s correlation between the two types of blood samples, 4 genes (*AOAH, ARG2, FKBP5* and *STAT3*) had Pearson’s R >0.94. The other gene *PAK2*, which had the lowest Pearson’s R (0.58), also had the lowest expression level relative to other genes (Figure 2C). These results indicated that NEC-associated DEGs could be detected in both whole blood and DBS.

### Stimulation of *S*.*epidermidis* increased ARG2

It remains a big challenge for neonatologists to differentiate NEC from sepsis, a type of neonatal infection that often co-exists with NEC but requires substantially different clinical management. Therefore at the end of this study, we examined whether the identified NEC-associated DEGs could be altered by infection. To do this, cord blood samples were stimulated *ex vivo* with *S*.*epidermidis* (SE) and lipopolysaccharides (LPS), representing infection caused by gram-positive and gram-negative bacteria independent of NEC lesions. After stimulation with SE and LPS, cord blood samples showed increased expression of cytokines (TNF-α and IL-10, by 3-14 fold), indicating an immune response to infection. Among the 5 target genes that were increased in severe NEC cases, *ARG2* was significantly increased by SE while *STAT3* tended to be increased by LPS (Figure 2D). This result suggested that the other 3 target genes (*AOAH, FKBP5* and *PAK2*) were associated with severe NEC and might be independent of infection.

## Discussion

Various NEC biomarkers have been studied to date. As blood remains the most readily available biofluid, in this proof-of-concept study, we used preterm pig model to address whether whole blood transcriptome could be used to identify novel NEC biomarker. The preterm pig is a valuable biomedical model for preterm infants due to their similarities in size, anatomy and clinical complications [25]. Similar to preterm infants, preterm pigs develop NEC spontaneously after formula feeding and show clinical symptoms such as lethargy, bloody stools and abdominal distension and discoloration that require discontinuation of feeding to avoid disease progression. During the first week of life, bloody stool is the only parameter that could predict NEC (mainly severe NEC) in pigs according to our retrospective analysis. However, bloody stool is not specific to NEC and could be attributed to milk allergy. Thus at this stage, NEC is suspicious and could only be diagnosed at necropsy. Whole blood transcriptome analyzed at such early stage of NEC revealed hundreds of DEGs that were mainly associated with severe NEC lesions, i.e., tissue necrosis and/or pneumatosis intestinalis. The NEC-associated DEGs were detectable in both whole blood and dried blood spots, and might differentiate NEC from infection. These results suggested the feasibility of using human blood transcriptome to develop early biomarkers for identifying infants who have severe NEC lesions. Such biomarkers will help avoid surgery on infants who can be treated medically, yet not delay operating on those who need to be treated surgically.

In this study, NEC-associated DEGs were discovered by transcriptome and target genes were validated by QPCR. Three genes (*AOAH, FKBP5* and *PAK2*) are noteworthy because they have been confirmed to increase in severe NEC while independent of infection, according to our *ex vivo* stimulation experiment. The expressions of these 3 genes were also highest in the two most severe NEC cases from discovery and validation cohort, respectively. Only these two pigs were diagnosed with score 6 in both small intestine and colon, showing extensive hemorrhage and necrotic lesions in both regions. Both pigs had bloody stools after 4 days of life, but clinical record and hematology results could not differentiate them from other NEC cases. This suggests that NEC-associated DEGs may be used to indicate NEC severity at early stage and may be superior to other clinical parameters. However, these target genes may not be optimal biomarkers for human infants, and further study is required to identify human specific biomarker. This could possibly be done using archived neonatal DBS samples, which are collected routinely across countries.

Biomarkers are usually associated with disease, but are not necessarily the cause of disease. However, some of our target genes worth further investigation on their roles in NEC pathogenesis. It has been hypothesized that TLR4 activation plays a crucial role in NEC development [26]. Rodent experimental model showed that TLR4 mediated Th17 lymphocyte influx, whereby release of IL-17 disrupted the intestinal barrier, leading to NEC. This process is dependent on STAT3, which drives Th17 differentiation [12]. Upon activation, lymphocytes in the mucosal immune system travel through mesenteric lymph nodes and thoracic duct, from where they circulate in the blood before reentering mucosal tissues. Thus, our observation of increased *STAT3* in the whole blood in severe NEC cases may reflect pathogenic lymphocytes influx to the gut. Moreover, one of our target genes *PAK2* was identified as a hub gene among all NEC-associated DEGs. This gene was suggested as a driver of colitis from a recent study of IBD, which has analyzed multiomics from mouse and human biopsies to generate the hypothesis [24]. Interestingly, this study indicated both *PAK2* and *STAT3* as key genes of IBD, but only *PAK2* was chosen to be validated as a therapeutic target. Inhibition of Pak signaling in animal model resolved bleeding and restored normal epithelial crypt morphology [24]. Whether *PAK2* also drives NEC development or whether targeting Pak signaling protects against NEC could be investigated in the future. Another gene that increased in severe NEC cases was *ARG2*, which encodes for arginase 2, a catabolizing enzyme of arginine. It has been shown that low arginine availability in the immature intestine is a risk factor of NEC, likely due to the result of poor intestinal perfusion [27]. Clinical trials in preterm infants also showed arginine supplementation prevented development of NEC [28]. Finally, *FKBP5* encodes for FK506 binding protein 5, which is a regulator of the glucocorticoid receptor (GR). Overexpression of *FKBP5* is known to result in glucocorticoid resistance, either by reducing affinity of GR for glucocorticoids or reducing nuclear translocation and activation of GR itself [29, 30]. As glucocorticoids are potent inhibitors of immune system, increased *FKBP5* in severe NEC cases may be related to inappropriate immune response.

In summary, we demonstrated that blood gene expression altered at early stage of NEC and candidate genes were closely related to severity of NEC lesions. Expressions of candidate genes could be used as early biomarkers to help decision making of treatment strategy. Further studies are encouraged to investigate how to combine multiple types of NEC biomarkers to provide better care.

## Acknowledgements

We thank Thomas Thymann, Anders Brunse, Duc Ninh Nguyen, Jing Sun, Kristine Holgersen, Jane Povlsen, Elin Skytte, Kristina Møller (University of Copenhagen) for their support to animal procedures and laboratory analyses.

## Author Contributions

XP analyzed and interpreted the transcriptome data. XP was the major contributor in writing the manuscript. TM and SR performed *ex vivo* stimulation experiment and analysis. FG and PS took part in the main study design and critically reviewed the manuscript. All authors read and approved the final manuscript.

## Data Availability

All sequencing and processed data will be deposited in the Gene Expression Omnibus (GEO).

## Reference

1. Patel, R.M., et al., Causes and timing of death in extremely premature infants from 2000 through 2011. N Engl J Med, 2015. 372(4): p. 331–40.

2. D’Angelo, G., et al., Current status of laboratory and imaging diagnosis of neonatal necrotizing enterocolitis. Ital J Pediatr, 2018. 44(1): p. 84.

3. Juhl, S.M., et al., Incidence and risk of necrotizing enterocolitis in Denmark from 1994-2014. PLoS One, 2019. 14(7): p. e0219268.

4. Luig, M., et al., Epidemiology of necrotizing enterocolitis--Part I: Changing regional trends in extremely preterm infants over 14 years. J Paediatr Child Health, 2005. 41(4): p. 169–73.

5. Ng, P.C., T.P. Ma, and H.S. Lam, The use of laboratory biomarkers for surveillance, diagnosis and prediction of clinical outcomes in neonatal sepsis and necrotising enterocolitis. Arch Dis Child Fetal Neonatal Ed, 2015. 100(5): p. F448–52.

6. Ng, E.W., et al., Gut-associated biomarkers L-FABP, I-FABP, and TFF3 and LIT score for diagnosis of surgical necrotizing enterocolitis in preterm infants. Ann Surg, 2013. 258(6): p. 1111–8.

7. Luo, J., et al., Early diagnosis of necrotizing enterocolitis by plasma RELMbeta and thrombocytopenia in preterm infants: A pilot study. Pediatr Neonatol, 2019. 60(4): p. 447–452.

8. Morrow, A.L., et al., Early microbial and metabolomic signatures predict later onset of necrotizing enterocolitis in preterm infants. Microbiome, 2013. 1(1): p. 13.

9. Ng, P.C., et al., Plasma miR-1290 Is a Novel and Specific Biomarker for Early Diagnosis of Necrotizing Enterocolitis-Biomarker Discovery with Prospective Cohort Evaluation. J Pediatr, 2019. 205: p. 83–90 e10.

10. Leaphart, C.L., et al., A critical role for TLR4 in the pathogenesis of necrotizing enterocolitis by modulating intestinal injury and repair. J Immunol, 2007. 179(7): p. 4808–20.

11. Sodhi, C.P., et al., Toll-like receptor-4 inhibits enterocyte proliferation via impaired beta-catenin signaling in necrotizing enterocolitis. Gastroenterology, 2010. 138(1): p. 185–96.

12. Egan, C.E., et al., Toll-like receptor 4-mediated lymphocyte influx induces neonatal necrotizing enterocolitis. J Clin Invest, 2016. 126(2): p. 495–508.

13. Cao, M., et al., Physical activity level is impaired and diet dependent in preterm newborn pigs. Pediatr Res, 2015. 78(2): p. 137–44.

14. Thymann, T., et al., Formula-feeding reduces lactose digestive capacity in neonatal pigs. Br J Nutr, 2006. 95(6): p. 1075–81.

15. Kim, D., et al., TopHat2: accurate alignment of transcriptomes in the presence of insertions, deletions and gene fusions. Genome Biol, 2013. 14(4): p. R36.

16. Anders, S., P.T. Pyl, and W. Huber, HTSeq--a Python framework to work with high-throughput sequencing data. Bioinformatics, 2015. 31(2): p. 166–9.

17. Love, M.I., W. Huber, and S. Anders, Moderated estimation of fold change and dispersion for RNA-seq data with DESeq2. Genome Biol, 2014. 15(12): p. 550.

18. Ye, J., et al., Primer-BLAST: a tool to design target-specific primers for polymerase chain reaction. BMC Bioinformatics, 2012. 13: p. 134.

19. Huang da, W., B.T. Sherman, and R.A. Lempicki, Bioinformatics enrichment tools: paths toward the comprehensive functional analysis of large gene lists. Nucleic Acids Res, 2009. 37(1): p. 1–13.

20. Liu, T., et al., Quantitative proteomic analysis of intracerebral hemorrhage in rats with a focus on brain energy metabolism. Brain Behav, 2018. 8(11): p. e01130.

21. Tsuchida, C., et al., Expression of REG family genes in human inflammatory bowel diseases and its regulation. Biochem Biophys Rep, 2017. 12: p. 198–205.

22. Bouzid, D., et al., Association of ZAP70 and PTPN6, but Not BANK1 or CLEC2D, with inflammatory bowel disease in the Tunisian population. Genet Test Mol Biomarkers, 2013. 17(4): p. 321–6.

23. Boros, E., et al., Elevated Expression of AXL May Contribute to the Epithelial-to-Mesenchymal Transition in Inflammatory Bowel Disease Patients. Mediators Inflamm, 2018. 2018: p. 3241406.

24. Lyons, J., et al., Integrated in vivo multiomics analysis identifies p21-activated kinase signaling as a driver of colitis. Sci Signal, 2018. 11(519).

25. Sangild, P.T., et al., Invited review: the preterm pig as a model in pediatric gastroenterology. J Anim Sci, 2013. 91(10): p. 4713–29.

26. Nino, D.F., C.P. Sodhi, and D.J. Hackam, Necrotizing enterocolitis: new insights into pathogenesis and mechanisms. Nat Rev Gastroenterol Hepatol, 2016. 13(10): p. 590–600.

27. Robinson, J.L., et al., Prematurity reduces citrulline-arginine-nitric oxide production and precedes the onset of necrotizing enterocolitis in piglets. Am J Physiol Gastrointest Liver Physiol, 2018. 315(4): p. G638–G649.

28. Shah, P.S., V.S. Shah, and L.E. Kelly, Arginine supplementation for prevention of necrotising enterocolitis in preterm infants. Cochrane Database Syst Rev, 2017. 4: p. CD004339.

29. Maltese, P., et al., Glucocorticoid resistance in Crohn’s disease and ulcerative colitis: an association study investigating GR and FKBP5 gene polymorphisms. Pharmacogenomics J, 2012. 12(5): p. 432–8.

30. Wochnik, G.M., et al., FK506-binding proteins 51 and 52 differentially regulate dynein interaction and nuclear translocation of the glucocorticoid receptor in mammalian cells. J Biol Chem, 2005. 280(6): p. 4609–16.

